# Human Handedness: Genetics, Microtubules, Neuropsychiatric Diseases and Brain Language Areas

**DOI:** 10.1101/454660

**Authors:** A. Wiberg, G. Douaud, M. Ng, Y. Al Omran, F. Alfaro-Almagro, J. Marchini, D.L. Bennett, S. Smith, D. Furniss

**Author notes:** These authors contributed equally to this work (A. Wiberg, G. Douaud).

## Abstract

**Background:** The skew in distribution of handedness is a uniquely human trait, and has fascinated researchers for centuries. The heritability of handedness is estimated at 25%, but defining genetic variants contributing to this trait has so far proved elusive.

**Methods:** We performed GWAS of self-reported handedness in UK Biobank, a prospective cohort study of ∼500,000 individuals. Furthermore, we investigated correlations between our associated SNPs and brain imaging-derived phenotypes (IDPs) from >9,000 individuals in UK Biobank, as well as between self-reported handedness and IDPs.

**Results:** Our association study of 38,322 left-handers vs 356,567 right-handers (excluding ambidextrous participants) revealed three genome-wide significant loci (rs199512, 17q21.31, p=4.1x10^−9^; rs45608532, 22q11.22, p=1.4x10^−8^; rs13017199, 2q34, p=3.3x10^−8^). In the imaging study, we found strong associations between rs199512 and diffusion MRI measures mainly in white matter tracts connecting language-related brain regions (p<2.0x10^−6^). Direct investigation between handedness and IDPs revealed numerous associations with functional connectivity between the same language-related areas of the brain. A second GWAS of non-right handers (n=44,631) vs right-handers (n=356,567) revealed an additional locus: rs3094128, 6p21.33, p=2.9x10^−8^. Three of the four associated loci (2q34, 17q21.31, 6p21.33) contain genes that encode microtubule-related proteins that are highly expressed in the brain: *MAP2, MAPT* and *TUBB*. These genes are strongly implicated in the pathogenesis of diseases that are known to affect an excess of left-handed people, including schizophrenia.

**Conclusions:** This is the first GWAS to identify genome-wide significant loci for human handedness in the general population, and the genes at these loci have biological plausibility in contributing to neurodevelopmental lateralization of brain organisation, which appears to predispose both to left-handedness and to certain neurodegenerative and psychiatric diseases.

## Introduction

The skew in distribution of handedness is a uniquely human trait, and has fascinated researchers for centuries. One of the most remarkable features of human motor control is that approximately 90% of the population has had a preference of using their right hand over the left since at least the Paleolithic period^1^. It is widely believed that the lateralisation of language in the left hemisphere accounts for the evolution of right-handedness in the majority of humans^2^. The extent to which such a population bias is under genetic influence has been a topic of considerable debate, but there are many features in support of a genetic basis for handedness. For example, left-handedness runs in families^3^, and concordance of handedness is greater in monozygotic twins than dizygotic twins, with an estimated heritability of 25%^4^. Early studies into the genetics of handedness assumed that a single gene locus with two or more alleles may determine handedness^5–7^, but with the advent of genome-wide association studies (GWAS), such genetic models of handedness determination have been abandoned in favour of a multilocus (i.e. polygenic) genetic model^8^.

Despite several GWAS, genome-wide significant loci for human handedness have thus far remained elusive^9,10^. Only one locus (*PCSK6* on chromosome 15) has been discovered in a series of studies measuring relative hand skill (rather than handedness *per se*) in a relatively small and specific cohort of individuals with reading disabilities^11,12^; this locus failed to replicate in the general population.

UK Biobank is a prospective cohort study of ∼500,000 participants aged between 40-69 who have undergone whole-genome genotyping at ∼800,000 SNPs, and have allowed linkage of these data with their medical records, lifestyle questionnaires, and physical and cognitive measures^13^. An imaging extension to the UK Biobank study includes brain imaging, which consists of six distinct modalities covering structural, diffusion and functional imaging^14^. We previously developed a fully automated image processing pipeline for UK Biobank raw imaging data (primarily based on the FMRIB Software Library, FSL^15^). This pipeline generates thousands of image-derived phenotypes (IDPs), which are distinct individual measures of brain structure and function, that can be used for genetic analysis^14,16,17,^. Using genotype, imaging and handedness data from UK Biobank, we aimed to discover correlations between: (1) genotype and handedness, (2) handedness related genotypes and IDPs, and (3) handedness phenotype and IDPs.

## Methods

### Genetic association study

We performed a GWAS to discover genetic loci associated with handedness using the UK Biobank resource. Participants were grouped by self-reported handedness as recorded in UK Biobank Data Field 1707 – participants could choose between “Right-handed”, “Left-handed”, and “Use both right and left hands equally” (ambidextrous).

Following sample- and SNP-based quality control (QC), we performed a GWAS of 38,322 left-handers against 356,567 right-handers of white British ancestry, excluding ambidextrous participants. We followed this with two further GWAS incorporating the ambidextrous participants: non-right-handers (n=44,631) vs right-handers (n=356,567); and left-handers (n=38,322) vs non-left-handers (n=362,866). Genome-wide association testing was undertaken across 547,011 common-frequency genotyped SNPs (minor allele frequency (MAF) ≥ 0.01) and ∼11 million imputed SNPs (MAF ≥ 0.01, Info Score ≥ 0.3) using a linear mixed non-infinitesimal model implemented in BOLT-LMM v. 2.3^18^ to account for population structure and relatedness. We assumed an additive genetic effect and conditioned on genotyping platform and genetic sex.

### *In Silico* Analyses

For the left- vs right-handers GWAS, we used FUMA^19^ to map genes to the associated loci, and quantified the expression of these genes across different tissue types. To gain insight into the biological pathways and gene sets that overlap with genes associated with handedness, we performed a gene-set analysis using MAGMA^20^, and a similar SNP-based enrichment analysis of the GWAS SNPs using XGR^21^. Brain expression quantitative trait loci (eQTLs) were obtained from the UK Brain Expression Consortium (UKBEC) dataset^22^ and from GTEx^23^ (See URLs). We performed LD score regression^24,25^ on summary-level statistics for the left- vs right-handers GWAS to estimate the SNP heritability, and to estimate the genetic correlation between handedness and various neurological and psychiatric diseases from publicly available summary-level GWAS data.

### Permutation analysis of microtubule associated genes

We created a list of all 348 genes associated with the gene ontology ‘Microtubule Cytoskeleton Organisation’ (GO:0000226) from the Broad Institute’s Gene Set Enrichment Analysis^26^ (see URLs). Using PLINK v1.9^27^, we pruned our 547,011 post-QC genotyped SNPs using an r^2^ threshold of <0.1, resulting in a set of 154,385 pruned proxy SNPs across the 22 autosomes. These proxy SNPs were randomly permuted 10,000 times into sets of four SNPs using a random number generator function in R v3.3.1, and for each set, we counted the number of times that at least one microtubule-associated gene from the ontology GO:0000226 was found within +/− 1Mb of each of the four SNPs in the set, based on the Gencode Release 28 (GRCh38.p12) gene set (see URLs).

### Imaging study of handedness genotype

We investigated the associations between each of the loci identified in the GWAS and the 3,144 IDPs available in almost 10,000 UK Biobank participants^14^. Then, to reveal the spatial extent of the relevant SNPs’ effects, we further analysed the brain images voxel-by-voxel using regression against the count of the non-reference allele (0, 1 and 2). Results were considered significant after Bonferroni correction for multiple comparisons (across IDPs and loci, or across voxels, respectively). Significant voxelwise, localised results were then used as starting points for the virtual reconstruction and identification of the white matter tracts to which they belong.

### Imaging study of handedness phenotype

Similarly, we carried out an analysis of self-reported handedness, excluding ambidextrous participants, in the subset of imaged UK Biobank participants across all 3,144 IDPs (n=721 left-handers, n=6,685 right-handers).

Full details of our methodology can be found in the **Supplementary Appendix**.

## Results

### Four novel genetic loci associated with human handedness

Comparing left-handers vs right-handers, we discovered three genome-wide significant associated loci: rs199512, 17q21.31, p=4.1x10^−9^; rs45608532, 22q11.22, p=1.4x10^−8^; rs13017199, 2q34, p=3.3x10^−8^. Comparing non-right-handers vs right-handers uncovered one further locus: rs3094128 at 6p21.33, p=2.9x10^−8^, and replicated the association at rs199512 on chromosome 17. Comparing left-handers vs non-left-handers, did not yield any new associated loci, but replicated the three loci from left- vs right-handers. For the four associated loci, we performed conditional analysis based on the top associated SNP at each locus, and did not observe any independent associations with handedness. The associated SNPs are summarised in **Table 1** and are displayed in **Figure 1**.

**Table 1.** **SNPs significantly associated with left-handedness**

| Chromosome | Position<sup>a</sup> | rsID | Effect Allele | EAf<sup>b</sup> | INFO<sup>c</sup> | OR (95% CI) | P | Candidate Genes |
| --- | --- | --- | --- | --- | --- | --- | --- | --- |
| 2 | 210246064 | rs13017199 | G | 0.369 | 0.997 | 1.04 (1.03-1.06) | 3.3 x 10<sup>-8</sup> | *MAP2* |
| 6* | 30694374 | rs3094128 | C | 0.219 | 1.000 | 0.95 (0.94-0.97) | 2.9 x 10<sup>-8</sup> | *TUBB* |
| 17 | 44857352 | rs199512 | C | 0.782 | 1.000 | 1.06 (1.04-1.07) | 4.1 x 10<sup>-9</sup> | *MAPT* |
| 22 | 23412190 | rs45608532 | A | 0.070 | 0.942 | 1.09 (1.06-1.12) | 1.4 x 10<sup>-8</sup> | *GNAZ, MICB* |

\*Only significant in non-right handers (i.e. left-handers + ambidextrous) vs right-handers GWAS
<sup>a</sup>Based on NCBI Genome Build 37 (hg19).
<sup>b</sup>The effect allele frequency in left handers (+/- ambidextrous depending on the analysis – see methods for details).
<sup>c</sup>The SNP INFO score for imputed SNPs

**Figure 1.**
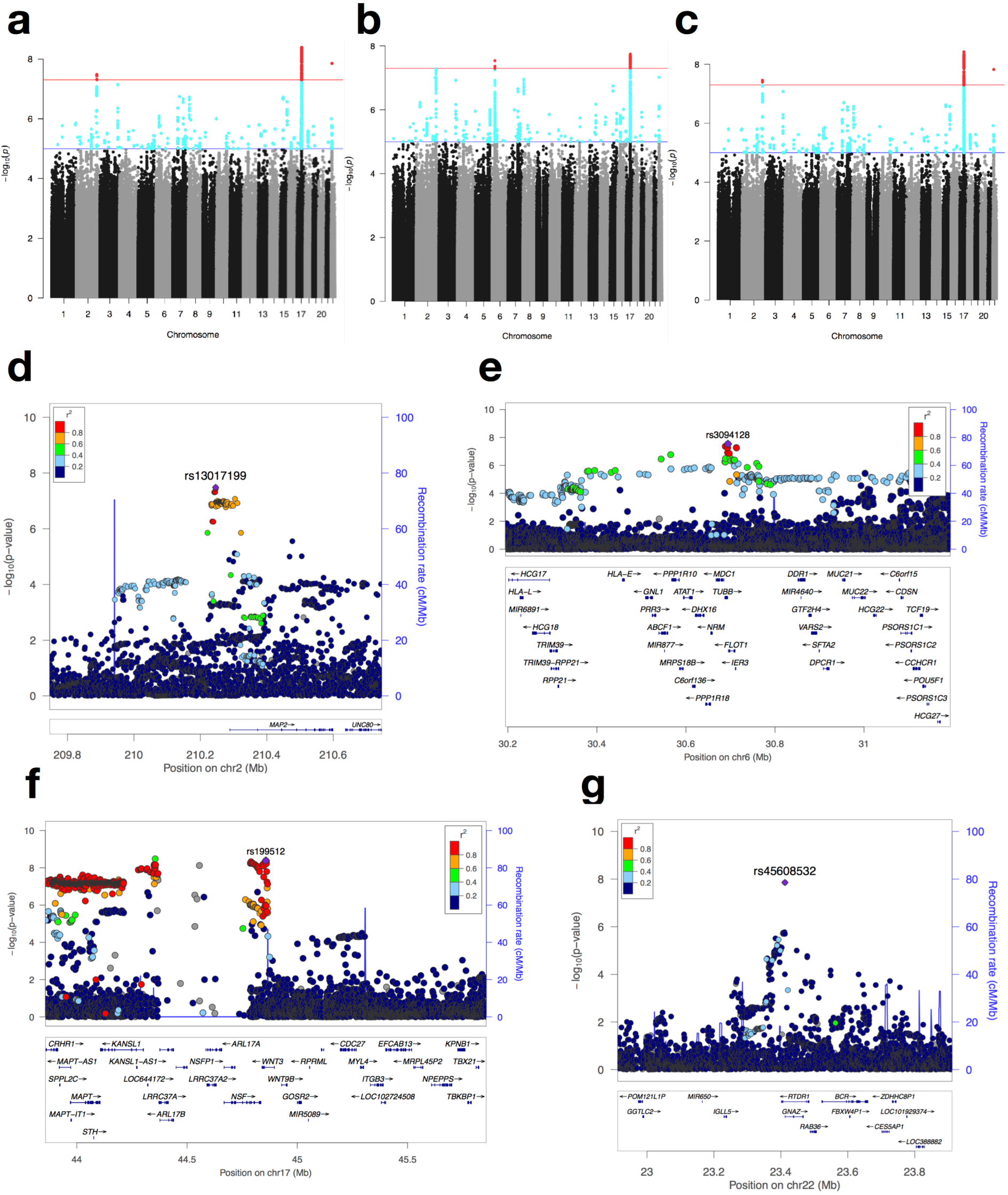
**Summary of GWAS of handedness. a-c**, Manhattan plots showing −log10 p-values for SNP associations in GWAS of: (**a**) right-handers vs left-handers (with ambidextrous participants excluded from cases and controls), (**b**) right-handers vs non-right-handers, and (**c**) left-handers vs non-left handers. The red line indicates the threshold for genome-wide significance (p=5x10^−8^), and variants coloured in red have genome-wide significant associations. **d-f**, Regional association plots of the associated SNPs at: (**d**) rs13017199 at 2q34, (**e**) rs3094128 at 6p21.33, (**f**) rs199512 at 17q21.31, and (**g**) rs45608532 at 22q11.22. The linkage-disequilibrium relationship between the lead SNP and the surrounding SNPs is colour-coded. The genomic positions of the SNPs and genes is based on Human Genome build hg19.

rs13017199 on chromosome 2 is ∼40kb upstream of *MAP2* - microtubule-associated protein 2. rs3094128 on chromosome 6 is approximately 1.2kb downstream of *TUBB* - Tubulin Beta Class 1. rs199512 on chromosome 17 is an intron within *WNT3*, and lies within a large linkage disequilibrium (LD) block (**Figure 1f**) within a common inversion polymorphism^28^. Notably, this region includes a further gene, *MAPT*, that encodes the microtubule associated protein tau; in fact, this region is commonly referred to as the *MAPT* locus. rs45608532 on chromosome 22 is an intronic variant within *RTDR1* - rhabdoid tumour deletion region gene 1 - and is upstream of *GNAZ* - a member of the G-protein family that is highly expressed in the brain^23^. Regional LocusZoom^29^ plots of the four associated SNPs are shown in **Figure 1d-1g**.

We sought to determine the empiric probability of three of four associated SNPs being within 1Mb of a gene affecting microtubule architecture occurring by chance alone. In only 299/10,000 random permutations, three or more of four SNPs are located within 1Mb of a microtubule-associated gene, giving a probability of <0.03 of this occurring by chance alone (**Supplementary Table S1**).

### Gene-set analysis and tissue expression

Positional gene mapping in FUMA^19^ of the left- vs right-handers GWAS summary statistics revealed a set of genes with high expression in various brain tissues (**Supplementary Figure S1**). We also performed a gene-set analysis on results from this GWAS using MAGMA^20^ - the four gene sets and gene ontology terms with the most overlapped genes pertained to neuronal morphogenesis, differentiation, migration and gliogenesis (**Supplementary Table S2**).

### Enrichment for neurodegenerative traits

Using XGR^21^, we implemented SNP-based enrichment analysis of 4,009 SNPs with p≤5x10^−5^ in the left- vs right-handers GWAS. The top two enrichments that we found were for Parkinson’s disease (p=2.6x10^−19^) and for ‘neurodegenerative disease’ (p=2.8x10^−12^); of the top eleven enrichments, nine were for neurological disorder phenotypes (**Supplementary Table S3).**

We performed LD score regression to examine correlations between handedness and neurodegenerative and psychiatric phenotypes. Our most statistically significant correlations were with Parkinson’s disease (r_g_=−0.2379, p=0.0071) and schizophrenia (r_g_=0.1324, p=0.0021) (**Supplementary Table S4**). Also using LD score regression, we calculated the heritability (h^2^) of handedness explained by all the SNPs in the left- vs right-handers GWAS to be 0.0121 (standard error 0.0014).

### Language-related brain structural connectivity is associated with handedness genotype

We found many significant genotype:IDP associations for rs199512, especially in white matter tracts using diffusion MRI (structural connectivity) measures (**Supplementary Table S5**). The two strongest associations were with the IDP labelled “TBSS superior longitudinal fasciculus R” (L1 measure, p=3.4x10^−9^) and “TBSS anterior limb of the internal capsule R” (L3 measure, p=3.0x10^−9^). These particular measures describe water diffusion along and perpendicular to the major tracts, respectively. Voxel-by-voxel analysis of the L1 maps showed bilateral localised effects (p<3.6x10^−7^) which were confirmed, using tractography, to be in the arcuate fasciculus/superior longitudinal fasciculus (SLF) III connecting language-related regions such as Broca’s area (Brodmann 44 and 45), as well as the planum temporale (**Figure 2A**). L3 maps revealed bilateral effects of the rs199512 polymorphism in white matter tracts connecting the anterior part of the supplementary motor area (pre-SMA) and in the uncinate fasciculus, as well as in the right arcuate fasciculus and cingulum bundle.

**Figure 2.**
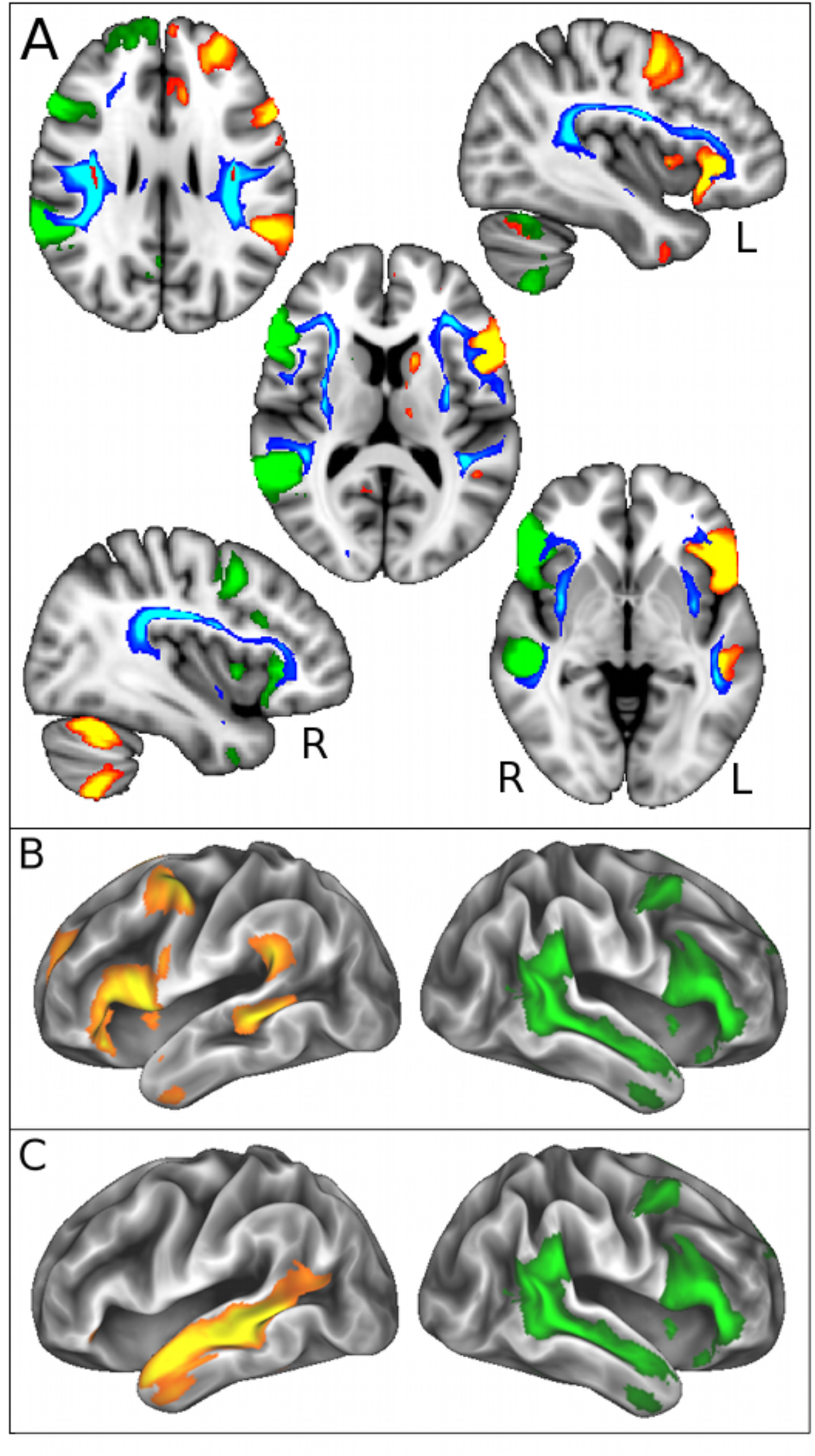
**White matter tracts associated with rs199512 connect language-related grey matter regions functionally involved with self-reported handedness. A.** Voxelwise effects in white matter associated with rs199512 (in red, p<3.6x10^−7^) were used as seeds for probabilistic tractography, which reconstructed the arcuate and superior longitudinal fasciculus (III) (in blue-light blue, thresholded for better visualisation at 250 samples). Results are overlaid on the MNI T1-weighted template (axial views: z = 27, 12, −3 mm; sagittal views: x = −39, 39 mm). These white matter tracts clearly link grey matter areas present in lateralised right- and left-sided language functional networks (in green and red-yellow, respectively, also shown in B). **B and C.** Left-handedness was most strongly associated with an increase in functional connectivity (temporal correlation) between right language functional network (in green, encompassing Broca’s areas, the planum temporale and superior temporal sulcus, Z>5), and a split of the left language functional network (in red-yellow, Broca’s areas and planum temporale shown in B., superior temporal sulcus shown in C., Z>5). These language-related functional networks are overlaid on the cortical surface.

### Language-related brain functional connectivity is associated with handedness

Directly comparing all 3,144 IDPs between left- and right-handers irrespective of genotype yielded numerous significant results, all but one using resting-state (functional connectivity) measures (**Supplementary Table S6A and S6B**). Remarkably, the top two hits demonstrated an *increase* of functional connectivity in left-handers between left-sided and right-sided language areas networks, including the same brain regions connected by white matter tracts discovered in our previous genotype and imaging analysis (**Figure 2A**). The strongest difference in functional connectivity between left- and right-handers was between a left-sided resting-state “language network” encompassing Broca’s areas (Brodmann 44 and 45) and the planum temporale, and its right-sided homolog (p=5.2x10^−44^) (**Figure 2B**). The second strongest association between handedness and functional connectivity was identified between the left superior temporal sulcus and the right-sided homolog to the language network (p=4.1x10^−31^) (**Figure 2C**). Only one significant IDP was a marker of structural connectivity in the white matter (“TBSS external capsule Left”, p=5.7x10^−6^, using the mode of anisotropy – “MO”). Voxelwise MO differences and tractography revealed higher structural connectivity in left-handers in the right corticospinal tract (corresponding to movement of the left part of the body), in the corpus callosum (which connects left and right hemispheres), and in the arcuate fasciculus, the same tract discovered in our genotype and imaging result.

**Supplementary Figure S2** summarises the key findings from the three arms of this study.

## Discussion

We have discovered four novel genetic loci associated with human handedness through genome-wide association in the UK Biobank cohort. Furthermore, we found statistically significant correlations between genotype at the most significant SNP, rs199512, and structural connectivity measures in white matter tracts connecting language-related brain areas, and strong associations between self-reported handedness and many functional connectivity measures, particularly in those same language regions (**Supplementary Figure S2**).

### Genetics and Handedness

Strikingly in the genetic association study, of the four significantly associated regions, three contained a gene encoding a microtubule component – *MAP2, TUBB* and *MAPT*. While further functional studies are required to prove causality between the statistically significant SNPs at our three loci and the three microtubule-related genes in question, we demonstrated through permutation analysis that this number of microtubule-related genes in proximity to our lead SNPs were unlikely to occur by chance alone. This suggests that these genes are part of a biological pathway that contributes to the phenotypic expression of non-right-handedness. Microtubules are an integral component of the neuronal cytoskeleton^30^, and particularly interesting in the context of phenotype such as handedness because of their key role in neuronal morphology, migration, transport and polarisation, and brain morphogenesis^31^.

*MAP2* is the gene that lies closest to rs13017199 on chromosome 2, also an eQTL of this gene. Aberrations in *MAP2* have been implicated in Parkinson’s disease^32^, and altered expression of *MAP2* has been reported within various brain structures including the hippocampus and prefrontal cortex in schizophrenia^33^. *TUBB*, as a tubulin gene, plays a crucial role in the mechanisms of CNS development including neuronal migration and axonal outgrowth and maintenance. Notably, a number of brain malformations have been associated with mutations in tubulin genes^34,35^. Of note, rs3094128, which lies 1.2kb from *TUBB*, is also an eQTL of *MICB*^23^, which is specifically expressed in the brain, and plays a crucial role in brain development and plasticity, and may mediate both genetic and environmental involvements in schizophrenia^36^. There is a plethora of literature linking left-handedness to an array of psychiatric disorders, particularly schizophrenia^37^. A meta-analysis of 50 studies concluded that non-right-handedness was significantly more common in schizophrenic participants (OR=1.84), and supports the view that there is a genetic link between handedness, brain lateralisation and schizophrenia^38–40^.

The most significantly associated SNP in the left- vs right-handers GWAS, rs199512, resides in a large LD block on chromosome 17 that contains *MAPT*, and is an eQTL of *MAPT* and *MAPT-AS1*^22,23^. There is a wealth of literature on variants in the *MAPT* gene that are associated with neurodegenerative phenotypes, which are collectively grouped under the name of tauopathies. Perhaps the best-known pathological associations of MAPT are Parkinson’s and Alzheimer’s diseases, with a recent study specifically demonstrating the genetic overlap between these two neurodegenerative disorders within this extended *MAPT* region^41^. Depositions of tau-associated neurofibrillary tangles have been shown to mediate neurodegeneration and clinical decline in Alzheimer’s disease and tau has thus been one of the favoured drug targets in this disorder^42^. Several polymorphisms in and around *MAPT* have been discovered in GWAS of Parkinson’s disease^43^. Those SNPs account for the genetic enrichment observed between handedness and Parkinson’s disease in both our SNP-based enrichment and LD score regression analyses.

### Imaging, Genotype and Handedness

Findings from previous neuroimaging studies of human handedness have been equivocal, except in the shape and depth of the central sulcus^44,45^, most likely owing to small- to medium-sized study populations^46,47^. The considerable size of the UK Biobank imaging cohort has allowed us to discover numerous novel correlations between handedness and imaging phenotypes.

In our genotype:imaging analysis, consistent with rs199512 being an eQTL of *MAPT* and *MAPT-AS1*, our top SNP yielded many highly significant associations with diffusion imaging IDPs, which characterise microstructural integrity in white matter tracts In particular, those differences were revealed most strongly in tracts linking Broca’s and temporoparietal junction areas (arcuate/SLF III). Language-related tracts such as the arcuate fasciculus have been consistently associated with schizophrenia and auditory hallucinations^48,49^, which might partially explain why our LD score regression analysis revealed correlations between our GWAS results and schizophrenia. The lack of lateralisation in our white matter results might be surprising at first, but seems to be in line with a recent study by ENIGMA, which failed to find any significant associations of handedness with grey matter asymmetries^50^. Strikingly however, these associated white matter tracts linked grey matter regions known to show the strongest asymmetries in the brain, from a very early developmental stage^51,52^. In addition, while it is not possible to formally associate the genetic contribution to handedness with these imaging results, it is notable that all of the grey matter regions connected by white matter tracts showing correlation with the rs199512 polymorphism make up the functional language networks that differ between left- and right-handers. This SNP therefore appears to have an anatomically and functionally relevant effect on brain connectivity, which lends credence to its putative role as one of the determinants of handedness.

The strongest IDP correlations of self-reported handedness were indeed found in functional connectivity *between* left- and right-sided language networks, with higher connectivity in left-handers. A stronger right dominance in left-handers could also be seen in lower functional connectivity between the right-homologous language network and the default-mode network, in other words a stronger suppression of the default-mode network to perhaps keep attention focused on language-relevant goals performed by these right-sided brain areas (p=1.9x10^−26^). While 96% of right-handers show a left-lateralisation of language function, about 15% of left-handers demonstrate bilateral function^53,54^. Our results, which reveal an increase of temporal correlation within subjects between left and right functional brain regions, suggest that more bilateral language function in left-handers might actually be a continuous trait. This might be mediated in part by the higher structural connectivity of the corpus callosum which left-handers also display.

In conclusion, this is the first GWAS to date to identify genome-wide significant loci for human handedness in the general population. The genes at these loci have biological plausibility in contributing to differences in neurodevelopmental connectivity of the language areas which might predispose both to left-handedness and certain neurodegenerative and psychiatric disorders. Furthermore, a direct effect of handedness was revealed in the lateralization of brain language function. Whether increased bilateral language function gives left-handers a cognitive advantage at verbal tasks remains to be investigated^54^. This study represents an important advance in our understanding of human handedness, and it offers some mechanistic insight into the observed correlation between handedness and certain neurodegenerative and psychiatric diseases.

## URLs

UK Biobank: www.ukbiobank.ac.uk; Gene Set Enrichment Analysis: http://software.broadinstitute.org/gsea/; LD Hub: http://ldsc.broadinstitute.org/ldhub/; XGR: http://galahad.well.ox.ac.uk:3020/; R: https://www.r-project.org; UK Brain Expression Consortium (UKBEC): http://www.braineac.org; Gencode: https://www.gencodegenes.org/releases/current.html/; FUMA: http://fuma.ctglab.nl/; MAGMA: https://ctg.cncr.nl/software/magma

